# GL261 glioblastoma induces delayed body weight gain and stunted skeletal muscle growth in young mice

**DOI:** 10.1101/2025.02.10.635159

**Authors:** Joshua R. Huot, Nicholas A. Jamnick, Fabrizio Pin, Patrick D. Livingston, Chandler S. Callaway, Andrea Bonetto

## Abstract

**Introduction:** The survival rate for children and adolescents has increased to over 85%. However, there is limited understanding of the impact of pediatric cancers on muscle development and physiology. Given that brain tumors alone account for 26% of all pediatric cancers, this study aimed to investigate the skeletal muscle consequences of tumor growth in young mice.

**Methods:** C2C12 myotubes were co-cultured with GL261 murine glioblastoma cells to assess myotube size. GL261 cells were then injected subcutaneously into 4-week-old male C57BL/6J mice. Animals were euthanized 28 days post-GL261 implantation. Muscle function was tested *in vivo* and *ex vivo*. Muscle protein synthesis was measured via the SUnSET method, and gene/protein expression levels were assessed via Western blotting and qPCR.

**Results:** *In vitro*, the C2C12 cultures exposed to GL261 exhibited myotube atrophy, consistent with a disrupted anabolic/catabolic balance. *In vivo*, carcass, heart, and fat mass were significantly reduced in the tumor-bearing mice. Skeletal muscle growth was impeded in the GL261 hosts, along with smaller muscle CSA. Both *in vivo* muscle torque and the *ex vivo* EDL muscle force were unchanged. At molecular level, the tumor hosts displayed reduced muscle protein synthesis and increased muscle protein ubiquitination, in disagreement with decreased muscle ubiquitin ligase mRNA expression.

**Conclusions:** Overall, we showed that GL261 tumors impact the growth of pediatric mice by stunting skeletal muscle development, decreasing muscle mass, reducing muscle fiber size, diminishing muscle protein synthesis, and altering protein catabolism signaling.

## INTRODUCTION

Eighty percent of adult cancer patients experience the hallmarks of ‘cachexia’, characterized by loss of appetite, reduced weight and skeletal muscle mass, increased fatigue, functional impairment and heightened chemotherapy toxicity. These symptoms ultimately lead to poor quality of life and reduced survival (1–3). Conversely, defining childhood cachexia is more challenging due to physiological differences related to puberty and age. Pubertal effects on height, weight, and body composition are also often overlooked in children with cancer (2). Additionally, inconsistent nutritional screenings and the lack of easily administered body composition assessments complicate the definition of cachexia in the pediatric cancer population, leading to its underutilization (4). Lastly, while many anticancer agents may reduce and prevent cachexia by counteracting tumor growth and progression, the pro-cachectic effects of cancer therapeutics in children remain largely unexplored. Of note, it is currently unclear which aspects of cachexia as induced by cancer or its treatments may be reversible or contribute to long-lasting metabolic defects. Consequently, conceptualizing pediatric cachexia has been difficult, despite its potential irreversible and long-term developmental consequences (4).

When the Survellance, Epidemiology, and End Results (SEER) program of the National Cancer Institute started collecting information on cancer statistics in 1975, 58% of children and 68% of adolescents were expected to survive their cancer diagnosis (5). Today, the survival rate for children and adolescents has increased to over 85% (6–8). This significant improvement can be attributed to advancements in treatments, surgical equipment, and access to medical care. Of all cancers, central nervous system (CNS) tumors are amongst the most common in pediatric patients, accounting for 26% of all cancer diagnoses (6, 9). Glioblastomas, in particular, are amongst the most common cancers in pediatric patients and make up approximately 15% of all CNS tumors in children (10). Generally, pediatric CNS cancers have more severe musculoskeletal consequences compared to hematological tumors, such as leukemias. Children with cancer often experience significant reductions in fat-free mass (11, 12) and caloric intake (13), along with muscle weakness (14, 15), neurological consequences (16), decreased quality of life, cognitive side effects (17), and heightened treatment-related toxicity (18). These factors collectively lead to poorer long-term health outcomes and higher all-cause mortality throughout their lives (19–21). Despite the well-documented short-term systemic effects and increases in all-cause mortality associated with childhood cancer, the molecular mechanisms driving these outcomes remain unclear.

Generally, tumor burden can drive skeletal muscle wasting by secreting pro-inflammatory cytokines that affect muscle, adipose and the CNS. These cytokines can lead to reduced anabolism, increased protein degradation, skeletal and cardiac muscle atrophy, decreased innervation, increased lipolysis, increased bone resorption, and behavioral and humoral changes in the brain (1, 22). Tumor- and host-derived cytokines such as insulin like growth factor (IGF)-1, tumor necrosis factor (TNF), and interleukin (IL)-6 are known to initiate skeletal muscle atrophy by inhibiting, among others, the PI3K/AKT/FOXO1/3 pathway, or by activating the STAT3, C/EBPβ and NF-κB signaling pathways, which in turn trigger the transcription of atrophy-related genes like Atrogin-1 and MuRF1, muscle specific ubiquitin ligases notoriously overactivated in association with muscle hypercatabolic states (22).

Research from ours and other laboratories has primarily focused on exploring the musculoskeletal consequences of colorectal or ovarian cancers, typical adult malignancies, using animal models employing older (>10 weeks of age) mice (23–27). Previous research using GL261 tumors examined the effects of chemotherapeutics on tumor growth and survival (28), as well as the impact of GL261 tumor mutations and therapeutics on cachexia (29). Although these studies reported reductions in body mass (-5% and -20%, respectively), they did not address the derangements in muscle homeostasis caused by the tumor. Additionally, the mice used were older than 7 weeks, beyond the typical window of the ‘pediatric’ mouse models. Despite the well described mechanisms of cancer cachexia driving skeletal muscle atrophy in adults, there is minimal understanding on the impact of pediatric cancer on muscle physiology and development.

In this study, we investigated the pro-cachexiogenic effects of GL261 glioblastoma tumors in young animals, aiming to recapitulate the effects of CNS tumors in the ‘pediatric’ population. Four-week-old male C57BL6/J were subcutaneously implanted with GL261 cells. Body weight and plantarflexion torque were assessed at baseline and at regular intervals. At the time of sacrifice, skeletal muscles, bones, and other tissues were collected, processed and stored for further analysis.

## MATERIALS & METHODS

### Cell culture

Murine GL261 cells were provided by Dr. Reza Saadzadeh (Indiana University School of Medicine, Indianapolis, IN, USA), cultured in high glucose Dulbecco’s modified Eagle’s medium (DMEM) medium (Cytiva, Marlborough, MA, USA) supplemented with 10% fetal bovine serum, 1% penicillin/streptomycin, and maintained in a 5% CO_2_, 37°C humidified incubator. Murine C2C12 skeletal myoblasts (ATCC) were grown in DMEM (10% fetal bovine serum and 1% penicillin/streptomycin) and maintained at 37°C in a 5% CO_2_ humidified atmosphere. Myotubes were generated by exposing the myoblasts to DMEM containing 2% horse serum and replacing the medium every other day for 5 days. To determine the effects of tumor-derived mediators on fiber size, myotubes were either exposed to 50% GL261-derived conditioned medium (CM), or co-cultured with GL261 cells growing in permeable inserts (Thermo Fisher Scientific, Waltham, MA, USA) for up to 48 hours.

### Generation of tumor-derived CM

Murine GL261 cells were grown in culture medium as described above. Once the cell layers reached >80% confluency, the culture medium was replaced with fresh medium containing 10% fetal bovine serum, 1% glutamine, 1% sodium pyruvate, and 1% penicillin and streptomycin. After 48 hours, the CM was collected, centrifuged at 1,200 rpm for 10 min at 4°C to remove cell debris, and stored at −80°C for further analyses.

### Assessment of myotube size

Cell layers were fixed in ice-cold acetone–methanol (50/50) and incubated with an anti-Myosin Heavy Chain antibody (MF-20, 1:200; Developmental Studies Hybridoma Bank, Iowa City, IA, USA) and an AlexaFluor 594-labelled secondary antibody (Invitrogen, Grand Island, NY, USA). Analysis of myotube size was performed by measuring the average diameter of long, multi-nucleated fibers (n = 250–350 per condition) avoiding regions of clustered nuclei on a calibrated image using the ImageJ 1.43 software (30), as in our previous works (26).

### Assessment of muscle cross sectional area (CSA)

To assess skeletal muscle atrophy, 10 μm-thick cryosections (CM1860, Leica Biosystems, Nussloch, Germany) of tibialis anterior muscles taken at the mid-belly were processed for immunostaining as described previously (31). Briefly, sections were blocked for 1 hour at room temperature and incubated overnight at 4°C with a dystrophin primary antibody [1:50, #MANDRA11(8B11), Developmental Studies Hybridoma Bank, Iowa City, IA, USA], followed by a 1 h secondary antibody (AlexaFluor 555, 1:1000, A21127, Thermo Fisher Scientific, MA, USA) incubation at room temperature. Entire dystrophin-stained sections were analyzed for CSA using a Lionheart LX automated microscope (BioTek Instruments, Winooski, VT, USA).

### Subcellular Fractionation

Nuclear and cytosolic proteins were extracted from a single gastrocnemius (∼125 mg.) muscle using modifications of a previously described protocol (32, 33). Muscle samples were thawed on ice in 1.5 mL of SEMH buffer (20 mM Hepes-KOH, pH 7.6; 220 mM Mannitol; 70 mM Sucrose; 1 mM EDTA) containing a protease/phosphatase inhibitor cocktail (#A32959 Thermo Fisher Scientific, Waltham, MA, USA). The sample was then homogenized using a drill with an attached Teflon pestle on ice at 1,000 rpm (2 sets x 20 strokes; 5 min between each set). The homogenate was transferred to a 1.5 mL microcentrifuge tube, placed on ice for 30 min, vortexed for 15 s, and was then centrifuged at 500 g for 10 min at 4LC. The supernatant (S1) was removed and placed into a 1.5 mL microcentrifuge tube and used to obtain the cytosolic and mitochondrial fractions. Pellet (P1) was resuspended in 1 mL of SEMH buffer and re-homogenized using a drill with an attached Teflon pestle on ice at 1,000 rpm (30 strokes re-suspended in 500 μL of SEMH buffer (with a protease/phosphatase inhibitor cocktail) and centrifuged at 1,000 g for 15 min at 4LC. The supernatant (S5) was discarded, and the pellet (P5) was resuspended in 500 μL of SEMH buffer (supplemented with protease/phosphatase inhibitor cocktail; 1:100) and centrifuged at 1,000 g for 15 min at 4LC. The supernatant (S6) was discarded, and the pellet (P6) was resuspended in 500 uL of SEMH buffer (supplemented with protease/phosphatase inhibitor cocktail) and centrifuged at 1,000 g for 15 min at 4LC. The supernatant (S7) was discarded and pellet (P7) was resuspended in 400 μL of NET buffer (20 mM Hepes, pH 7.9; 1.5 mM MgCl2; 1.5 M NaCl; 0.2 EDTA; 20% Glycerol; 1% Triton-X-100) containing a protease/phosphatase inhibitor cocktail (1:100), kept on ice for 30 min (vortexed every 10 min for 15 s), then passed 10 times through a 2.5-mL syringe and a 20- gauge needle, followed by a freeze thaw, then passage 10 times through a 22-gauge needle, followed by a final centrifugation at 9,000 g for 30 min at 4LC. The supernatant (S8) was the resultant nuclear fraction and pellet (P8) was discarded. To generate the cytosolic fraction, the supernatant (S1) was centrifuged at 2,000 g for 15 min at 4LC, transferred to a 1.5 mL microcentrifuge tube, and the pellet (P2) was discarded. The supernatant (S2) was centrifuged at 10,000 g for 30 min at 4LC and the supernatant (S3) was transferred to a 1.5 mL microcentrifuge tube, resulting in the cytosolic fraction. The pellet (P3) was resuspended in SEMH buffer, centrifuged at 12,000 g for 30 min at 4LC and supernatant (S4) was discarded. The pellet (P4) was resuspended in Sucrose buffer (10 mM HEPES, pH=7.6; 0.5 M Sucrose; 2 mM PMSF) supplemented with inhibitors, resulting in the mitochondrial fraction (stored at -80LC for future use). Purity of the subcellular fractionations was determined using antibodies for histone 3 (H3) (nuclei) and lactate dehydrogenase (LDH) (cytosol).

### Western blotting

Total protein extracts were obtained by lysing cell layers or homogenizing 100 mg quadriceps muscle tissue in radioimmunoprecipitation assay (RIPA) buffer (150 mM NaCl, 1.0% NP-40, 0.5% sodium deoxycholate, 0.1% sodium dodecyl sulfate (SDS), and 50 mM Tris, pH 8.0) completed with protease (Roche, Indianapolis, IN, USA) and phosphatase (Thermo Scientific, Waltham, MA, USA) inhibitor cocktails. Cell debris were removed by centrifugation (15 min, 14 000 g), and the supernatant was collected and stored at −80°C. Protein concentration was determined using the bicinchoninic acid (BCA) protein assay method (Thermo Scientific, Waltham, MA, USA). Protein extracts (15-40 mg) were then electrophoresed in 4–15% gradient SDS Criterion Tris-HCl precast gels (Bio-Rad, Hercules, CA, USA). Gels were transferred to nitrocellulose membranes (Bio-Rad, Hercules, CA, USA). Membranes were blocked with 8% BSA in TBS-Tween (0.1%) at room temperature for 1 h, followed by an overnight incubation with diluted antibody in 8% BSA in TBS-Tween (0.02% Sodium Azide) at 4°C with gentle shaking. After washing with TBST-T the membrane was incubated at room temperature for 1 h with either Anti-rabbit IgG (H + L) DyLight 800 or Anti-mouse IgG (H + L) DyLight 600 Secondary (Cell Signaling Technologies, Danvers, MA, USA). Blots were then visualized with Odyssey Infrared Imaging System (LI-COR Biosciences, Lincoln, NE, USA). Optical density measurements were taken using the ImageJ Analyzer software. Antibodies used were pSTAT1 (#9167), STAT1 (#9712), pSTAT3^Y705^ (#9145), STAT3 (#8768), pAkt^S473^ (#4060), Akt (#9272), pP38 MAPK (T180/Y182) (#4511), P38 MAPK (#9212), pERK1/2 (#4370), ERK1/2 (#4695), p4EBP1 (#2855), 4EBP1 (#9644), VDAC (#4866), pFoxO3a^Ser253^ (#9466), FoxO3a (#12829), OPA1 (#80471), pmTOR^Ser2448^ (#5536), mTOR (#2983), H3 (#4499), LDH (#2012), Cytochrome C (#11940), Tom20 (#42406), Mitofusin 2 (#9482) and Ubiquitin (#3933) from Cell Signaling Technologies, α-Tubulin (#12G10) from Developmental Studies Hybridoma Bank, LC3BI/II (#L7643) from Sigma, PGC-1α (AB3242) from EMD Millipore, puromycin (EQ0001) from Kerafast, and COX-IV (ab14744) from Abcam. Primary antibody dilution was 1:1000.

### Real-time quantitative polymerase chain reaction (qRT-PCR)

RNA from quadriceps was isolated using the miRNeasy Mini kit (Qiagen, Valencia, CA, USA) and following the protocol provided by the manufacturer. RNA was quantified by using a Synergy H1 spectrophotometer (BioTek, Winooski, VT, USA). Total RNA was reverse transcribed to cDNA by using the Verso cDNA kit (Thermo Fisher Scientific, Waltham, MA, USA). Transcript levels were measured by Real-Time PCR (Light Cycler 96, Roche) taking advantage of the TaqMan gene expression assay system (Life Technologies, Carlsbad, CA, USA). Expression levels for IGF-1 (Mm00439560), Atrogin-1 (Mm00499523_m1), MuRF-1 (Mm01185221_m1), and MUSA1 (Mm00505343_m1) were detected using Taqman Probes. Gene expression was normalized to -binding protein (TBP) (Mm01277042_m1) levels using the standard 2 methods.

### Animals

All experiments were conducted with the approval of the Institutional Animal Care and Use Committee at Indiana University School of Medicine and were in compliance with the National Institutes of Health Guidelines for Use and care of Laboratory Animals. In order to investigate the effect of murine glioblastomas on skeletal muscle mass in young animals, 4- week-old C57BL/6J (The Jackson Laboratory, Bar Harbor, ME, USA) male mice (n= 15) were subcutaneously injected with GL261 cells (1.0 x 10^6^) and monitored daily. Control mice received an equal volume of vehicle injected subcutaneously (n= 5). Mice were weighed daily then euthanized under isoflurane anesthesia after 28 days. Tibialis anterior muscles were frozen in liquid nitrogen cooled isopentane for histology. The remaining tissues were harvested and weighed, then snap frozen in liquid nitrogen and stored at −80°C for further studies.

### Protein Synthesis

For protein synthesis measurements, the established and validated SUnSET method using puromycin was used as previously described (34, 35). Briefly, 30 min prior to sacrifice, animals were treated with 0.04 μmol/g body mass puromycin (P-1033, A.G. Scientific, San Diego, CA, USA) intraperitoneally. Puromycin incorporation was determined via Western blot analysis as described below.

### *In Vivo* Muscle Contractility

Animals underwent *in vivo* plantarflexion torque assessment (Aurora Scientific Inc, Aurora, Canada), as previously described (36). Briefly, the left hind foot was taped to the force transducer and positioned to where the foot and tibia were aligned at 90°. The knee was then clamped at the femoral condyles, avoiding compression of the fibular nerve. Two disposable monopolar electrodes (Natus Neurology, Middleton, WI, USA) were placed subcutaneously posterior/medial to the knee to stimulate the tibial nerve. Maximum twitch torque was first determined using supramaximal stimulations (0.2 ms square wave pulse). Peak plantarflexion torque was then assessed following a supramaximal square wave stimulation (0.2 ms) delivered at 125Hz stimulation frequency (37).

### *Ex vivo* muscle contractility

EDL muscles were subjected to whole-muscle contractility assessment, as done previously (38). The EDLs were dissected, and stainless-steel hooks were tied to both tendons using 4-0 silk sutures. The muscles were placed between a mounted force transducer (Aurora Scientific Inc, Aurora, Canada) and incubated in a stimulation bath containing Tyrode solution (121 mM NaCl, 5.0 mM KCl, 1.8 mM CaCl_2_, 0.5 mM MgCl_2_, 0.4 mM NaH_2_PO_4_, 24 mM NaHCO_3_, 0.1 mM EDTA, and 5.5 mM glucose) supplemented with continuous O_2_/CO_2_ (95/5%). Following a 10-minute incubation period, the maximum twitch force was obtained by determining optimal muscle length (L0) and force-frequency relationships were assessed using a supramaximal incremental frequency stimulation sequence (10, 25, 40, 60, 80, 100, 125 and 150 Hz for 350 ms). Force data was collected and analyzed with the Dynamic Muscle Control/Data Acquisition and Dynamic Muscle Control Data Analysis programs (Aurora Scientific Inc, Aurora, Canada) and EDL muscle weight and L0 were used to determine specific force.

### *In Vivo* Electrophysiology

Triceps *surae* muscles of all animals were subjected to electrophysiological functional assessment using the Sierra Summit 3–12 Channel EMG (Cadwell Laboratories Incorporated, Kennewick, WA), as performed previously (39). Two 28- gauge stimulating needle electrodes (Natus Neurology, Middleton, WI) were used to stimulate the sciatic nerve of the left hindlimb, a duo shielded ring electrode was used for recording, and a ground electrode was placed over the animal’s tail. Baseline-to-peak and peak-to-peak compound muscle action potential (CMAP) responses were recorded utilizing supramaximal stimulations (constant current intensity: <10 mA; pulse duration: 0.1 ms). In addition to CMAP responses, all animals were assessed for peak to-peak single motor unit potential (SMUP) responses, using the incremental stimulation technique, as previously described (37, 39). Briefly, the sciatic nerve was sub maximally stimulated until stable, minimal all-or-none responses occurred. Ten successive SMUP increments were recorded and averaged. Baseline- to peak CMAP amplitudes were used for comparison between experimental groups, and motor unit number estimation (MUNE) was calculated using peak-to-peak CMAP and average SMUP, using the following equation: MUNE = CMAP amplitude (peak-to-peak)/average SMUP (peak- to-peak).

### Enzyme-linked immunosorbent assay

IL-6 and IGF-1 ELISA’s were performed using Quantikine (Minneapolis, MN, USA) ELISA Kit (IGF-1, #MG100; IL-6, #M6000B-1]. Briefly, 50 μL of diluent was added to each well, followed by 50 μL of sample (plasma for IGF-1 ELISA was diluted 1:500), standard, and control. Plates were incubated at room temperature for 2 hours. Plates were then aspirated, washed (5x) and 100 μL of conjugate was added to each well, plates were incubated at room temperature for another 2 hours. Wells were aspirated/washed (5x) and 100 μL of substrate solution was added to each well, plates were incubated at room temperature for 30 mins. Following incubation, 100 μL of stop solution was added to each well. Optical density was determined using a microplate reader set to 450 nm. Concentrations were calculated from standard curve. Samples were run in duplicate.

### Statistics

Statistical analyses were performed using GraphPad Prism 9.4.1 (GraphPad Software, San Diego, CA, USA). Two-tailed Student’s t-tests were employed to determine differences between control and GL261 hosts. A 2-way repeated-measures analysis of variance (ANOVA) was performed, followed by Bonferroni’s post hoc comparisons, for longitudinal measures and ex vivo muscle contractility of the EDL. In general, variance was tested throughout our study, and in most cases, there were no significant differences. When a variance was significant, a Welch’s test was used. Statistical significance was set at p <0.05 and data was presented as means ± SD, unless otherwise noted.

## RESULTS

### GL261 cells promote myotube thinning and upregulation of pro-atrophic signaling pathways

To examine the influence of glioblastoma cells on myotube diameter and cell signaling, we co-cultured C2C12 myotubes with GL261 cells using permeable inserts. After a 48-hour co-culture, we observed a significant reduction in myotube diameter (-35% *vs.* control; CON: 17.4±0.5 µm, GL261: 11.3±1.6 µm; p<0.001) **(Figure 1 A-B)**. Similarly, C2C12 cultures exposed to GL261 CM displayed significant myotube atrophy **(Figure S1 A-B)**. The atrophic phenotype in the C2C12 myotubes co-cultured with the GL261 cells was also accompanied by a significant upregulation of pERK (p<0.001; +252% vs. control), p-p38 (p<0.001; +125% vs. control), and pSTAT1 (p<0.01; +73% vs. control). Conversely, p-mTOR was significantly downregulated (p<0.05; -18% vs. control), with no changes observed in p-4EBP1, pAKT, pSTAT3^Tyr705^ and total ubiquitin levels **(Figure 2 A-H)**.

**Figure 1.**
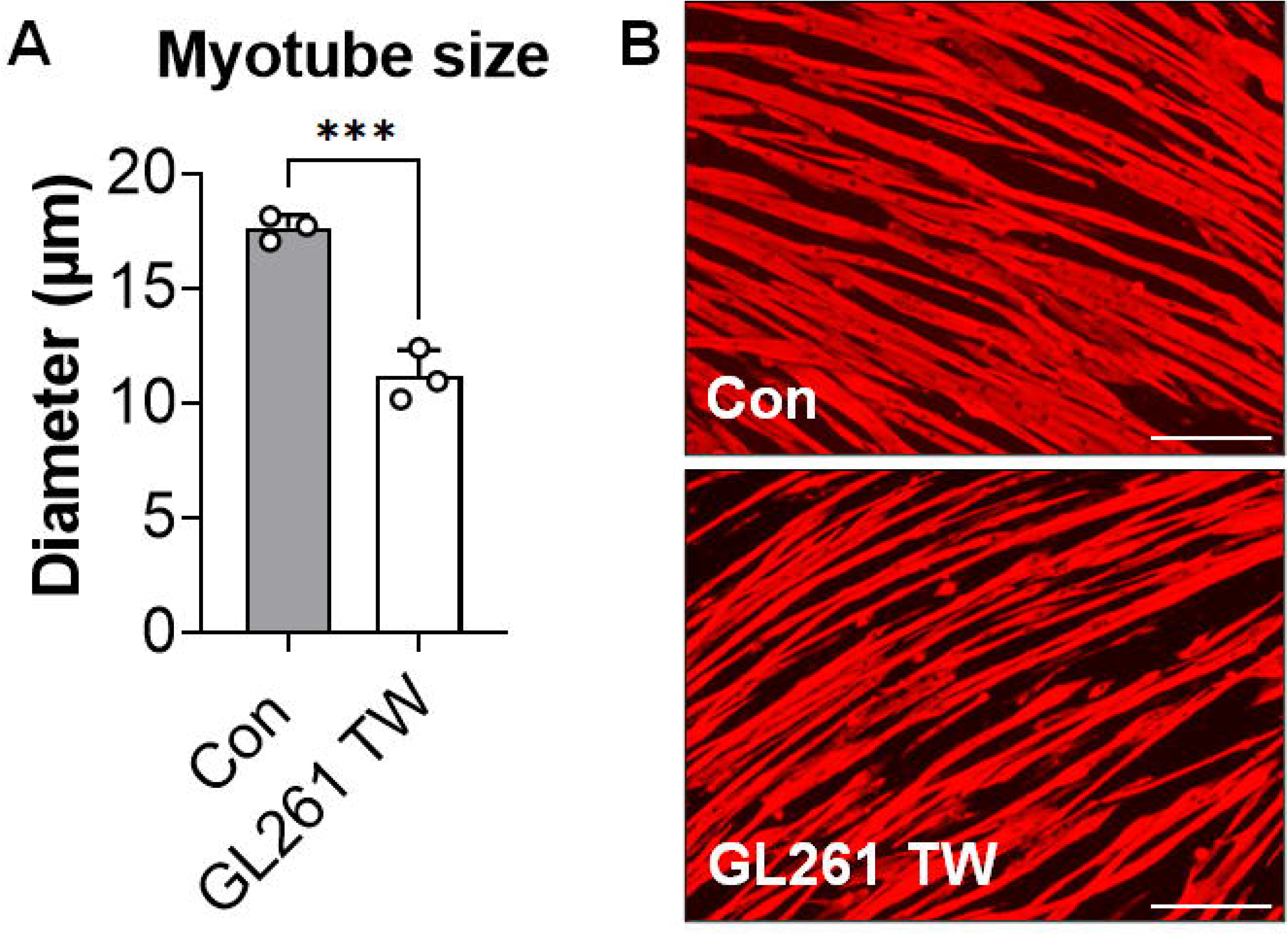
GL261 cells promote myotube atrophy. Myotube diameter (expressed as µm) of C2C12 myotubes co-cultured with GL261 cells **(A)**. Representative images of control (Con) and myotubes co-cultured with GL261 cells via permeable inserts (Transwells, TW) **(B)**. Scale bar: 100 µm. Data expressed as mean ± SD. Significant differences: ***p<0.001.

**Figure 2.**
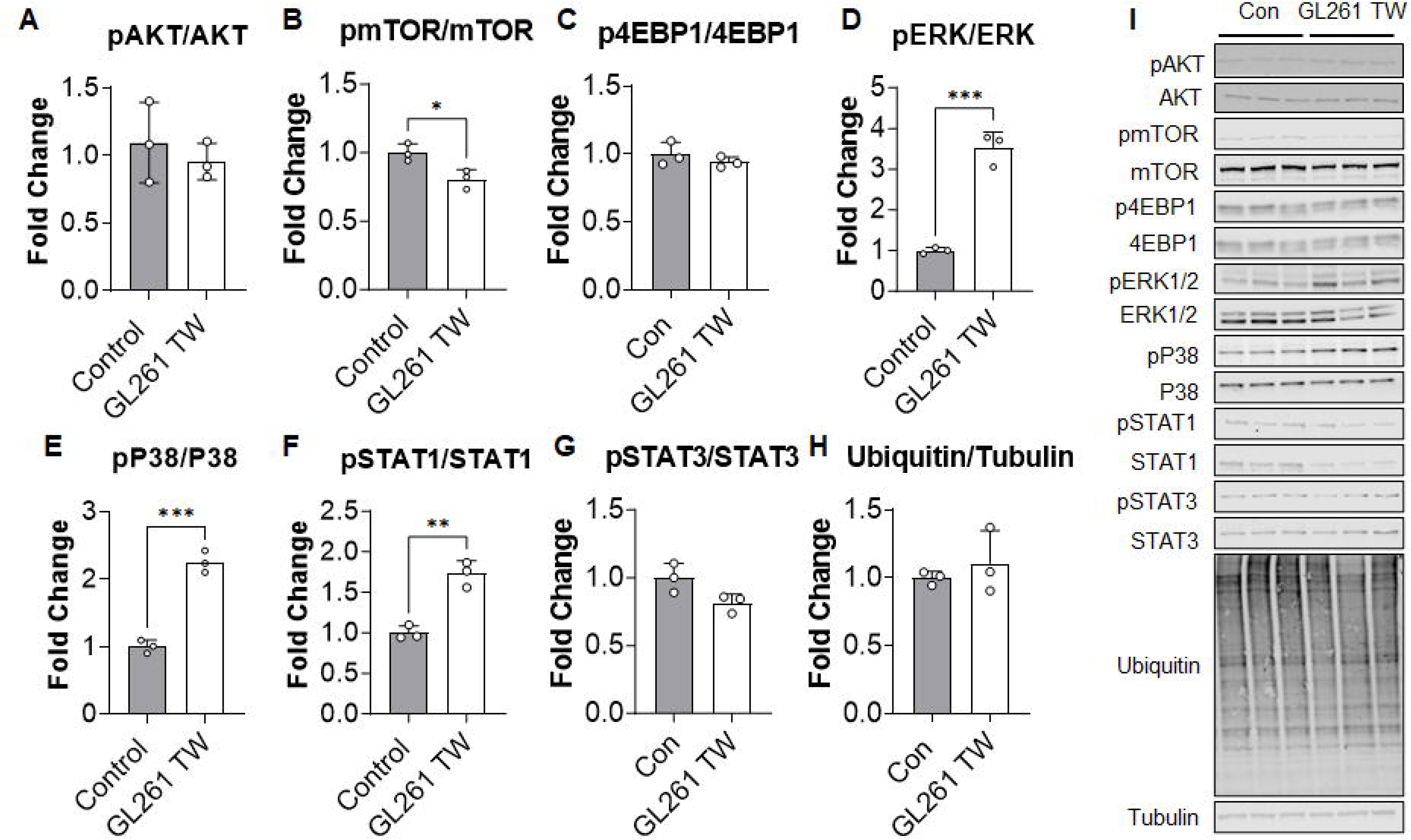
GL261-derived factors disrupt the anabolic/catabolic balance in C2C12 myotubes. Protein expression (assessed via Western blotting) of pAK/AKT **(A)**, pmTOR/mTOR **(B)**, p4EBP1/4EBP1 **(C)**, pERK/ERK **(D)**, pP38/P38 **(E)**, pSTAT1/STAT1 **(F)**, pSTAT3/STAT3 **(G)**, and ubiquitin **(H)** in C2C12 cells co-cultured with GL261 cells via permeable inserts (Transwells). Representative western blottings **(I)**. Data expressed as fold-change (mean ± SD). Significant differences: *p<0.05, **p<0.01, ***p<0.001.

### GL261 hosts exhibit stunted growth and changes in organ masses

To assess the impact of GL261 on the musculoskeletal development of young mice, male C57BL/6J mice were subcutaneously injected with 1.0 x 10^6^ cancer cells **(Figure S2)**. As shown by the body weight curves in **Figure 3A**, the tumors grew progressively larger reaching an average of 3.0 ± 1.5 g at time of euthanasia (28 days after tumor implantation) **(Figure 3B)**. This tumor growth significantly affected the overall body growth, as indicated by a reduced tumor-free carcass weight (-10% *vs.* control) in the GL261 group (CON: 62.4±3.8; GL261: 56.4±4.0; p<0.05) (**Figure 3C**). Interestingly, the GL261 hosts presented diminished gonadal fat accumulation (p<0.0001; -46% *vs.* control) (**Figure S3A**) and smaller heart mass (p<0.05; -7% *vs.* control) (**Figure S3B**), while the spleen size was significantly increased (p<0.001; +114% *vs.* control) (**Figure S3C**). There was a significant increase in liver mass (p<0.05: +15% *vs.* control) **(Figure S3D)**, whereas no changes in kidney size were observed **(Figure S3E)**.

**Figure 3.**
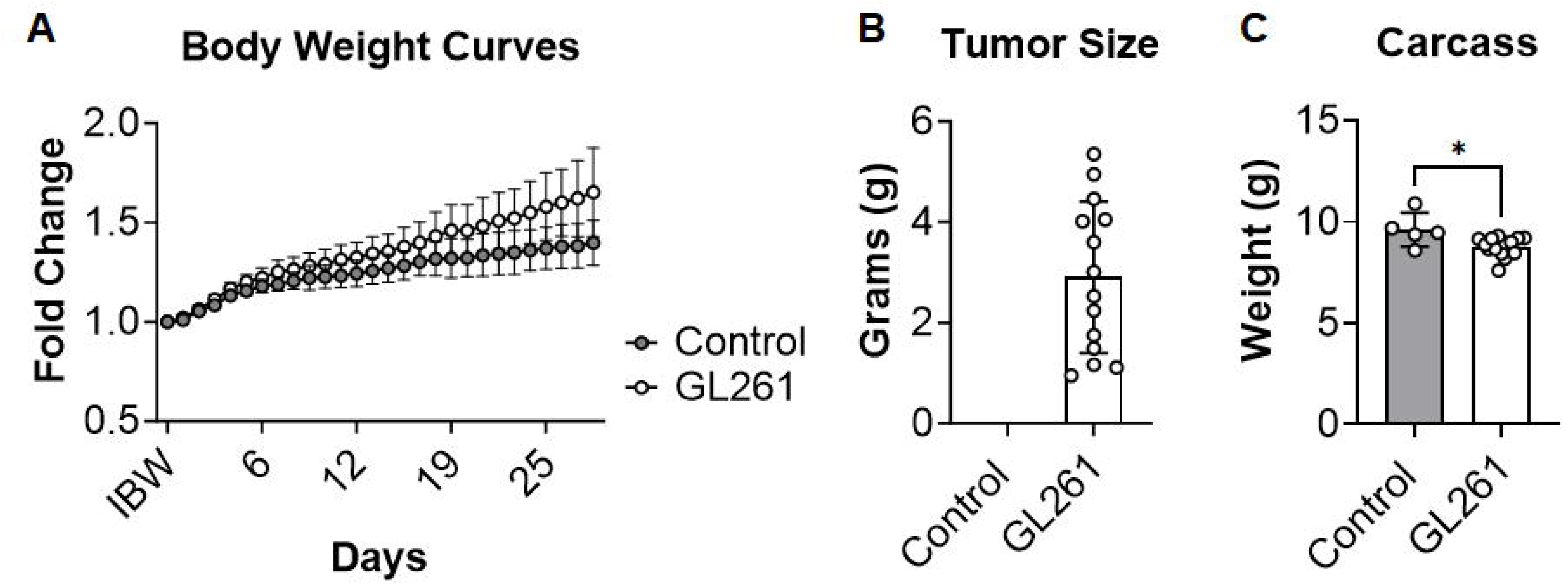
GL261 Tumors mitigate gains in body weight and diminished carcass weight. Body weight change (*vs.* initial body weight (IBW; expressed as fold-change) in control mice and GL261 hosts **(A)**. Tumor size (expressed in grams; g) **(B)** and carcass weights (normalized to initial body weight) **(C)**. Data expressed as mean ± SD. Significant differences: *p<0.05.

### Skeletal muscle size is diminished in the GL261 hosts

The GL261 hosts displayed stunted muscle growth compared to the controls. In particular, the quadriceps (p<0.01; -16% *vs.* control) and tibialis anterior muscles (p<0.05; -8% *vs*. control) were significantly smaller in the tumor hosts compared to the controls **(Figure 4A-B)**, while no statistical difference was found in the gastrocnemius mass **(Figure 4C)**. Correspondingly, the quantification of the muscle CSA revealed a reduction in average muscle fiber size in the tibialis anterior muscle of GL261 hosts (-17% *vs.* control; CON: 1502±183 µm^2^, GL261 1252±222 µm^2^; p<0.05) (**Figure 4D**).

**Figure 4.**
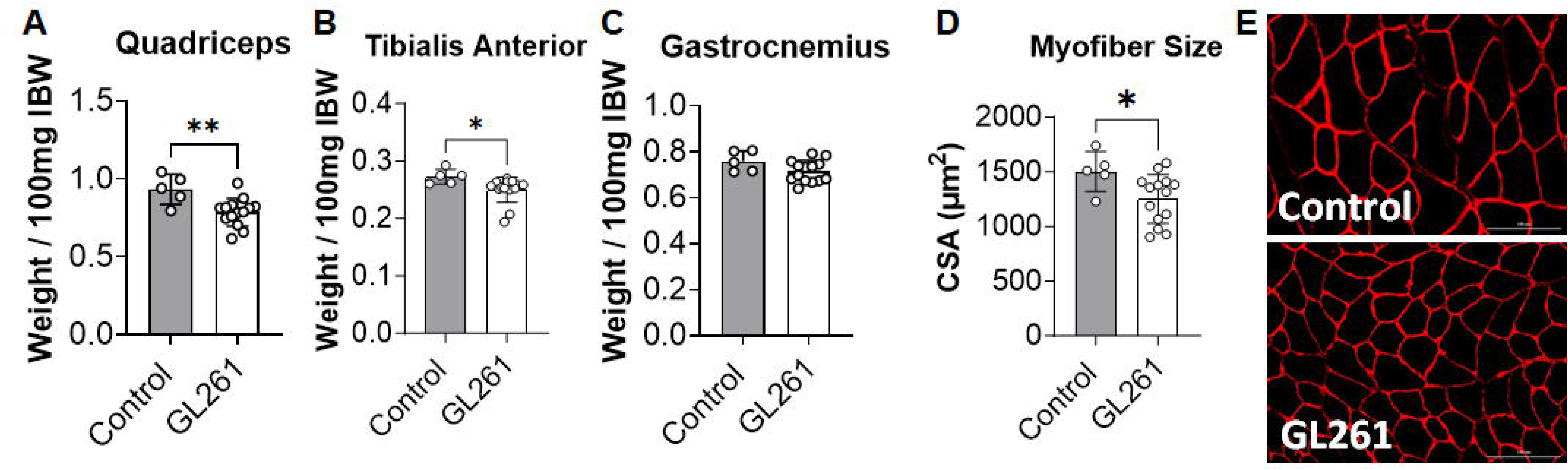
GL261 tumors reduce skeletal muscle weight and myofiber cross sectional area. Quadriceps **(A)**, tibialis anterior **(B)** and gastrocnemius **(C)** muscle weights from control and GL261-bearing mice were normalized to IBW and expressed as weight/100 mg IBW. Average cross-sectional area (CSA) of myofibers was expressed as µm^2^ **(D)**. Representative image of myofiber cross sectional area for control and GL261 hosts. Scale bar: 100 µm **(E)**. Data expressed as mean ± SD. Significant differences: *p<0.05, **p<0.01.

### GL261 hosts exhibited reductions in *ex vivo* muscular force and alterations in neuromuscular function

Isolated EDL muscles from GL261 hosts displayed a significant reduction in muscular force (p<0.01) compared to the controls **(Figure S4A-B)**, whereas the plantarflexion muscle torque was unchanged **(Figure S4C)**. Since we previously showed that impaired motor neuron function, denervation, and disrupted neuromuscular junction (NMJs) occur in cachexia (40), we then sought to investigate functional indices of motor unit connectivity also in the mice bearing GL261 tumors. The electrophysiological assessment of GL261 hosts revealed unchanged baseline-to-peak CMAP compared to the control mice **(Figure S4D)**. Consistent with our previous observations, we found significant increases in SMUP values in the tumor hosts (p<0.05; +54% *vs.* controls) **(Figure S4E)**, also consistent with reduced MUNE, suggestive of a decrease in motor unit numbers (p<0.05; -33% vs. controls) **(Figure S4F).**

### GL261 hosts exhibit abnormal skeletal muscle signaling

Similar to what is typically observed in older tumor-bearing mice, also in the young tumor hosts there were stark differences in the cell signaling cascades associated with onset of skeletal muscle atrophy **(Figure 5)**. Interestingly, we found a significant upregulation of pAKT in the quadriceps muscle of GL261 bearers compared to the healthy mice (p<0.05; +342% *vs.* control) **(Figure 5A)**. Consistent with previous data (41–43), we observed an increase in the LC3BII/I ratio (p<0.05; +84 *vs.* control) **(Figure 5B)**, suggesting that the autophagy-dependent catabolism was overactivated in the skeletal muscle of GL261 hosts. Additionally, there was a significant upregulation of STAT3 phosphorylation (p<0.05; +66% *vs.* control) in the nuclear subcellular fraction **(Figure 5E)**, in line with elevated circulating IL-6 levels **(Figure S5A)**. The levels of pFoxO3a^Ser253^, an upstream regulator of E3 ubiquitin ligase expression (44, 45), were found increased in the muscle cytosolic fraction (p<0.05; +211 vs. control) **(Figure 5D)**. Looking at regulators of mitochondrial homeostasis (in whole quadriceps muscle), we found that expression of PGC1α, a prominent regulator of mitochondrial biogenesis, was increased in the GL261 hosts (p<0.05; +98% *vs.* control) (46) **(Figure 5C)**, whereas the expression of VDAC (p<0.05; +40 *vs.* control), often associated with the activation of mitophagy (47, 48) **(Figure 6A)**, and mitofusin 2 (p<0.05; +47 vs. control), a marker of mitochondrial fusion (49) **(Figure 6C)**, were increased in the muscle of the tumor-bearing mice. No differences were observed in the expression of Tom20, OPA1, Cytochrome C, and COXIV proteins **(Figure 6B, D-F)**.

**Figure 5.**
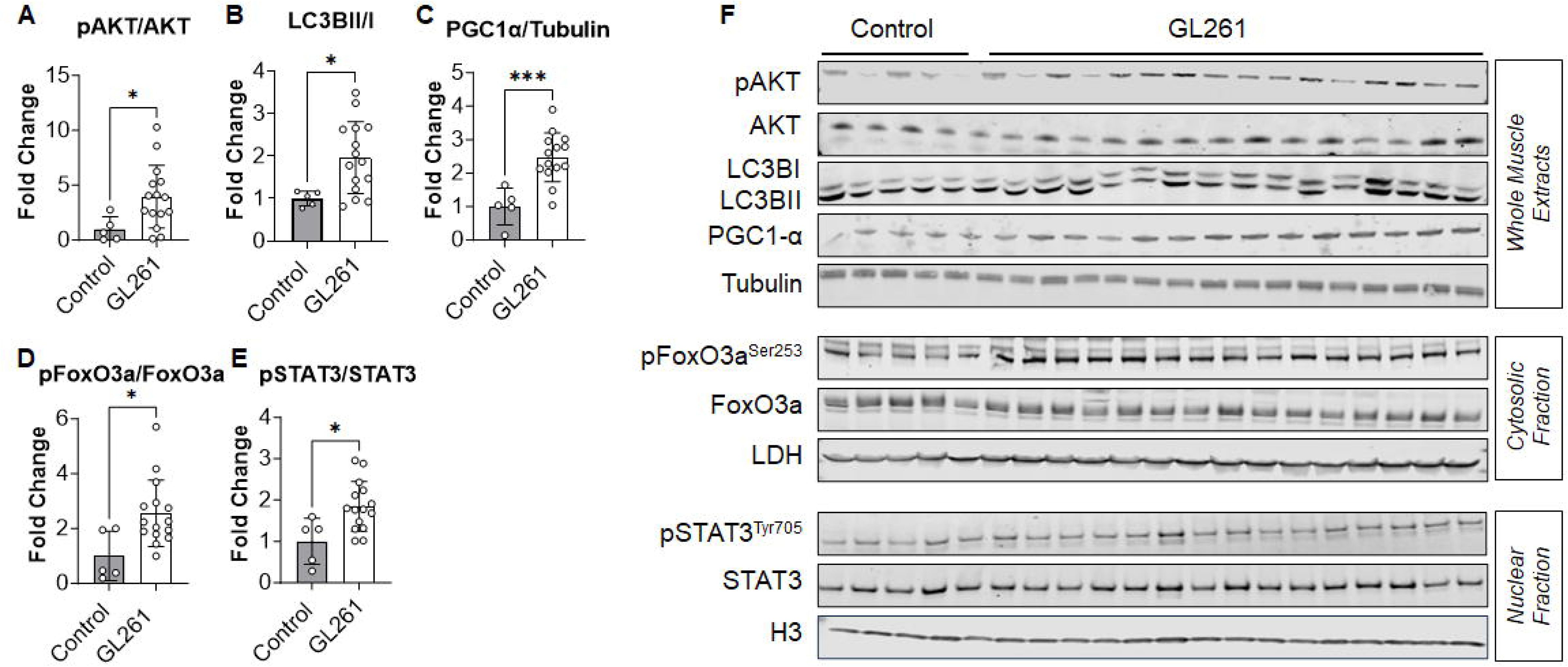
GL261 hosts display increased cell signaling associated with autophagy and muscle wasting. Protein expression (via Western blotting) of pAKT **(A)**, LC3BII/I ratio **(B)**, PGC-1α **(C)**, nuclear pSTAT3^Tyr705^ **(D)**, cytosolic pFoxO3a^Ser253^ **(E)** in the muscle of controls and GL261 hosts. Representative western blottings **(F)**. Data expressed as fold-change (mean ± SD). Significant differences: *p<0.05, **p<0.01.

**Figure 6.**
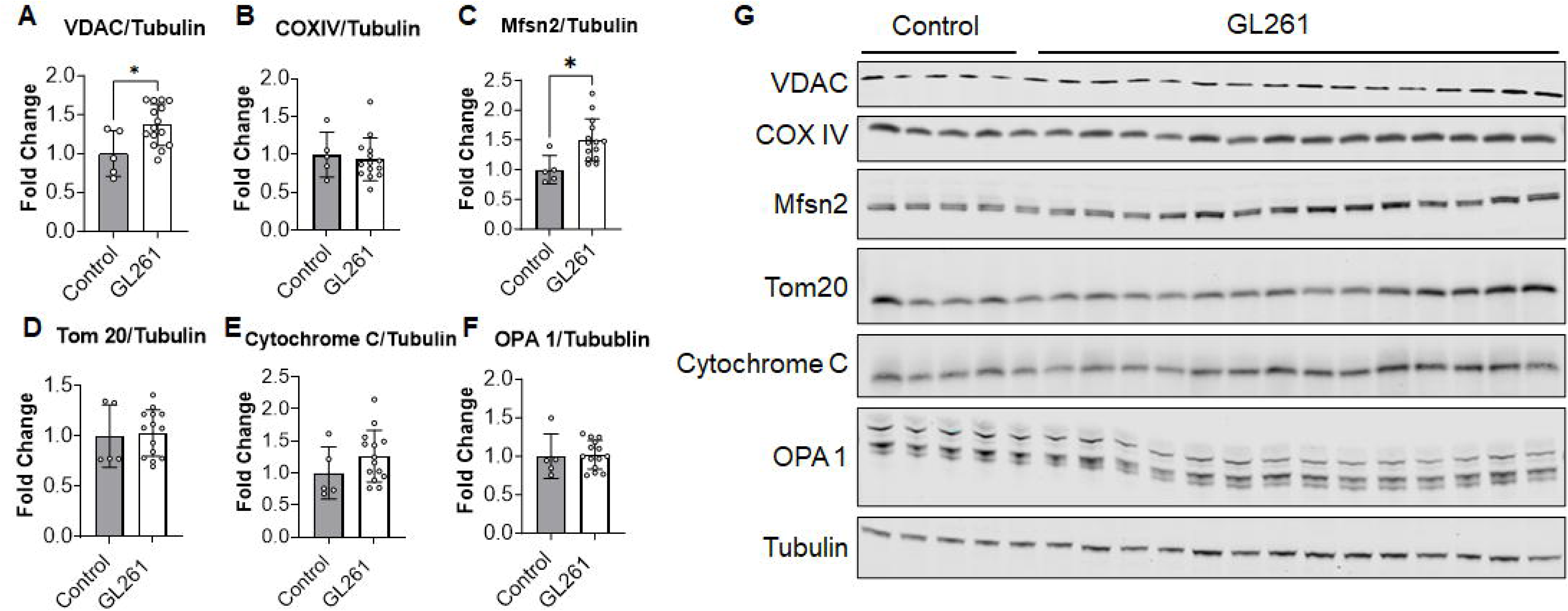
GL261 hosts exhibit abnormal mitophagy and mitochondrial turnover. Protein expression (via Western blotting) of VDAC **(A)**, COXIV **(B)**, Mitofusin 2 (Msfn2) **(C)**, Tom 20 **(D)**, Cytochrome C **(E)**, and OPA1 **(F)** in the muscle of controls and GL261 hosts. Representative western blottings **(G)**. Data expressed as fold-change (mean ± SD).Significant differences: *p<0.05.

In line with previous data reported in adult mouse models for the study of cancer cachexia (50, 51), mice carrying the GL261 tumors also exhibited increased ubiquitination (p<0.01; +54% *vs.* control), indicating enhanced protein degradation **(Figure 7A, C)**. To further investigate the mechanisms associated with skeletal muscle turnover, we assessed protein synthesis by determining the puromycin incorporation in the skeletal muscle of control animals and tumor hosts and found that protein synthesis was significantly reduced in the GL261 hosts (p<0.05; - 30% vs. control; **Figure 7B, C**). Along with evidence that IGF-1 gene expression was significantly downregulated in the muscle of GL261 hosts (p<0.05; -22% vs. control) **(Figure 7D)**, these observations suggest that protein homeostasis was severely dysregulated in the muscle of mice carrying glioblastomas. Surprisingly, the expression of muscle-specific E3 ubiquitin ligases, which are normally upregulated in conditions of cancer cachexia, was markedly downregulated in the GL261 hosts (Atrogin-1: p<0.05; -38% *vs.* control; MuRF-1: p<0.01; -54% *vs.* control; MUSA1: p<0.05; -58% *vs.* control) **(Figure 7E-G)**.

**Figure 7.**
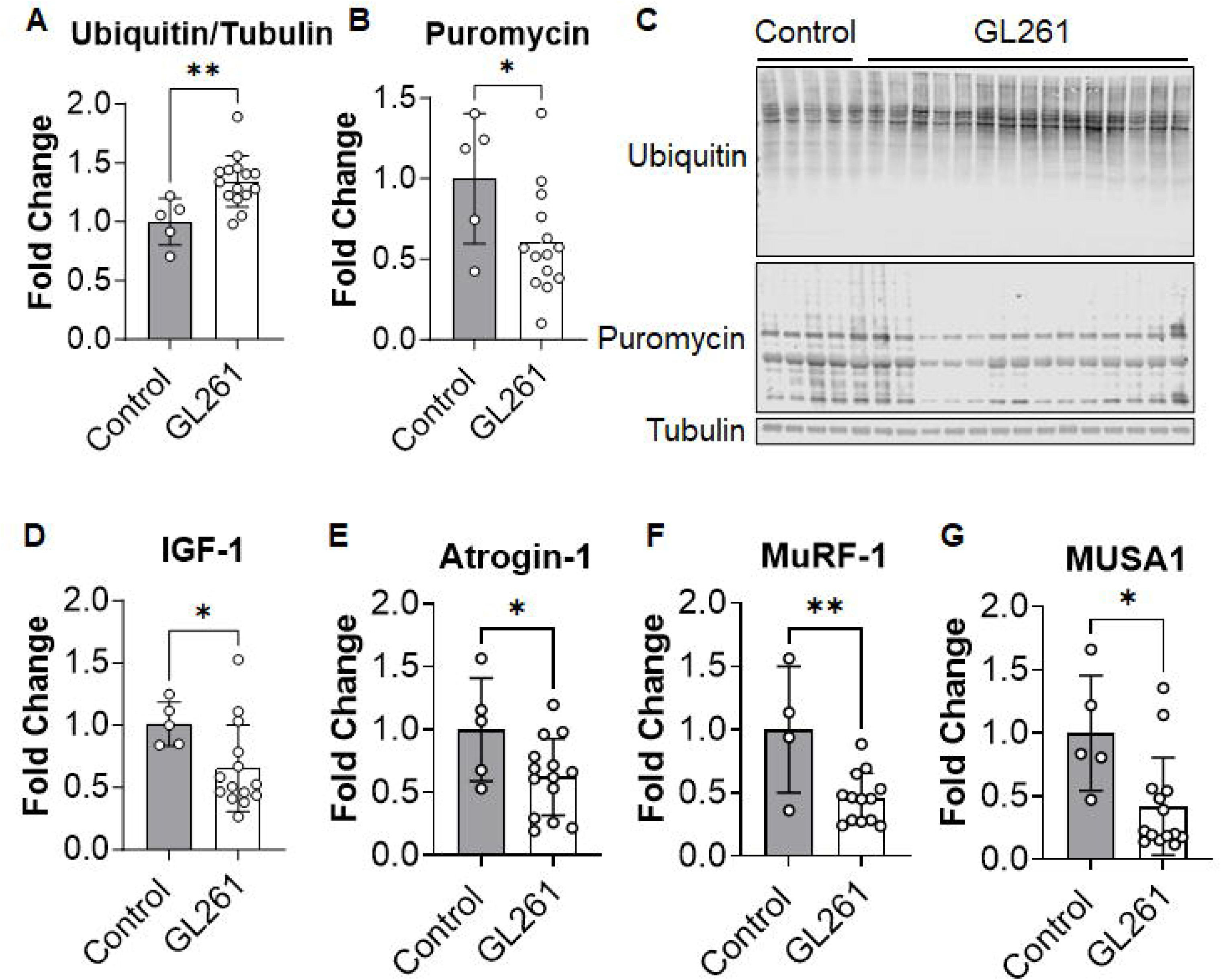
GL261 hosts display reduced muscle protein synthesis, increased protein degradation, and reduced expression of muscle-specific ubiquitin ligases. Protein expression (via Western blotting) of ubiquitin **(A)** and puromycin incorporation **(B)** in quadriceps muscle of control and GL261 hosts. Representative western blottings **(C)**. Tubulin was used as loading control. Data expressed as fold-change (mean ± SD). Gene expression (via qPCR) of IGF-1 **(D)**, Atrogin1 **(E)**, MuRF-1 **(F)**, and MUSA1 **(G)** in the quadriceps muscle of controls and GL261 hosts. Gene expression expressed as fold-change vs. control and reported as mean ± SD. Significant differences: *p<0.05, **p<0.01.

## DISCUSSION

Cancer-induced cachexia is a multi-organ wasting syndrome, mainly characterized by loss of skeletal muscle mass and leading to muscle weakness and worsened survival in patients (3, 52, 53). By employing adult (>8 weeks of age) mouse models of cancer, including colorectal (23, 24, 54–56), pancreatic (57, 58) and ovarian (27, 59), we and others have demonstrated that tumor burden triggers the onset of cachexia and that the drivers of cancer-induced skeletal muscle wasting may include reduced muscle anabolism, increased catabolism, muscle denervation, and decreased mitochondrial function (3, 23, 57, 59–63). Altogether, these abnormalities typically lead to damaging effects throughout the body (52, 64, 65).

Whether this is also true in young, pediatric tumor hosts is currently under investigation and the mechanisms that govern the overall stunted development due to childhood cancer diagnoses remain to be elucidated. Given that childhood cancer survivorship has increased over the past decades due to advances in screening and treatment, it has become increasingly urgent to understand these cellular mechanisms (4, 7, 8). Here, we aimed to examine the effects of GL261 glioblastoma tumors in young mice by focusing on changes in body composition, muscle force and skeletal muscle molecular signaling. To our knowledge, no studies have previously examined the cancer-associated impact on the growth and function of the musculoskeletal system in young/pediatric animals (<5 weeks of age).

Our *in vitro* experiments demonstrated that GL261 glioblastoma cells promote myotube atrophy and upregulate cell signaling markers associated with muscle wasting. This is consistent with previous research demonstrating that myotubes exposed to conditioned media from glioma cells (i.e., CHG5 and U87) exhibit altered protein synthesis, although the effects on myotube size were not assessed (66). Here, we found higher phosphorylation levels of ERK1/2, p38 and STAT1 in C2C12 myotubes co-cultured with GL261 cells. This data is in agreement with previous observations showing that increased activation of ERK1/2 MAPK was associated with reduced myotube size, whereas counteracting the ERK1/2 signaling was reported as a potential tool to promote muscle growth (67) and/or to prevent muscle wasting in cancer settings (68). Another MAPK, p38, was previously found to phosphorylate *Ser12* on p300 to stimulate C/EBPβ acetylation (69), ultimately leading to enhanced proteolysis through activated ubiquitin genes and autophagy pathways (70–72). Myotube thinning and reduced force generation were also associated with increased STAT1 phosphorylation, mainly due to high levels of interferon gamma (IFN-γ) (73), a condition frequently observed during aging. Lastly, we reported that reduced mTOR signaling was evident in myotubes co-cultured with GL261 cells, consistent with prior experimental conditions associated with reduced muscle protein synthesis and decreased myotube diameter (74–78). Altogether, these findings demonstrate that GL261 cells can promote C2C12 myotube shrinkage via activation of a typical pro-atrophic cell signaling cascade, thereby suggesting a potential pro-cachexiogenic effect of glioblastoma tumors.

To further investigate this point, we implanted young (4-week-old) mice with GL261 tumors. Our results demonstrate that GL261 glioblastomas delayed body growth and lowered skeletal muscle mass compared to normal, tumor-free mice. These observations align with previous research demonstrating large tumor growth and evidence of body weight loss in rodents implanted with GL261 cells (28, 29, 79). Although one study did not report the age of the mice and specifics of weight loss (79), another reported a reduction in body weight of ∼6%, which was further exacerbated by chemotherapy (i.e., vincristine and mebendazole) (28). While the most recent research reported a decrease in body weight of 20% (29), it is noteworthy that none of these studies described any derangements in muscle homeostasis. Interestingly, Cui and colleagues reported that implantation of glioma cells in adult, 8-week-old rodents caused severe cachexia, accompanied by complex metabolic derangements (66), further corroborating the possibility that the onset of brain cancers may promote musculoskeletal defects also in younger hosts. In our study, employing young animals, we demonstrated that the GL261 hosts exhibit modest losses of tumor-free body weight, also accompanied by decreases in carcass weights and stunted growth. Of note, pediatric patients affected with cancer frequently present mild or no body mass loss (12, 80, 81), along with reductions in fat free mass (12, 14, 15, 82, 83), thus suggesting that our model is representative of the clinical scenario.

In adult cancer cachexia mouse models, we generally observe a reduction in skeletal muscle size as well as a decrease in muscle strength (24, 26, 27). Whether these are a direct consequence of tumor growth also in pediatric models remains to be fully understood. In the current study we detected significant depletion of skeletal muscles in the tumor-bearing mice compared to the controls. Notably, our data align with clinical observations of reductions in lean muscle mass and weakness in pediatric cancer patients (12, 15, 84–87) and childhood cancer survivors (11, 12, 14, 15). Although we did not observe a reduction in *in vivo* plantarflexion torque, we did observe a reduction in *ex vivo* force in the EDL muscle that aligns with previous data from older mouse cancer models (24–27) and pediatric clinical data (14, 88–90), as well as from our previous study conducted in pediatric animals exposed to chemotherapy (91). Our research group has previously reported that cancer and chemotherapy can promote disruptions in neuromuscular function and the onset of a denervation-like phenotype (91), likely contributing to muscle atrophy and weakness (25). Here, we observed increased SMUP along with drastically reduced MUNE, a measure associated with muscle weakness, in the GL261 hosts. Therefore, our data suggests that the GL261 bearers present a negative effect on skeletal muscle weight, muscular force and neuromuscular function.

The pathophysiological mechanisms behind skeletal muscle wasting in cancer have been extensively studied using traditional pre-clinical mouse models employing adult animals. To explore whether similar mechanisms also drive the stunted development observed in young rodents carrying GL261 tumors, we then investigated the activation of signaling pathways known to contribute to skeletal muscle atrophy in cancer cachexia. Notably, we found elevated levels of nuclear p-STAT3^Tyr705^, consistent with increased circulating IL-6, which is well known to promote muscle wasting in cancer (73, 92, 93). Additionally, GL261-bearing mice showed increments in muscle protein ubiquitination, suggesting high protein turnover consequential to the activation of the ubiquitin proteasome system (94–97), as well as elevated LC3BII/I ratio, indicative of enhanced autophagy (98).

Skeletal muscle wasting, as occurring during different pathological states and during aging, is frequently accompanied by elevated expression of markers associated with the ubiquitin proteasome system, including the E3 muscle-specific ubiquitin ligases (i.e., MUSA1, MuRF-1, and Atrogin 1). An intriguing finding in the present study was the lower expression of MUSA1, MuRF-1, and Atrogin 1 in the muscle of GL261 hosts. Given that the purported mechanisms of action during hypercatabolism and cachexia is usually the nuclear translocation of FoxOs, which increases E3 muscle-specific ubiquitin ligase transcription (44, 99–102), our observation that p- FoxO3^Ser253^ levels are higher in the cytosolic subcellular fraction in the muscle of GL261 hosts, also consistent with increased levels of the upstream modulator p-AKT^Ser473^ (103–107), would seem to be in line with the diminished E3 ligase expression (22, 44, 101, 108). Interestingly, despite the sharp contrast of our observations with the abundance of previous data showing that upregulation of E3 muscle-specific ubiquitin ligases is a hallmark of cachectic mouse muscles (24, 27, 44, 45, 109–111), a previous study also observed a downregulation of E3 ligases in sarcopenic aged mice (30 months of age) which was attributed to mitigated FoxO transcription via increased p-AKT and IGF-1R signaling (112). Nonetheless, the similar findings observed in the present study of depressed E3 ligase expression modulated by FoxO localization are noteworthy, especially considering the differing age and disease states of these mice. Furthermore, we also observed reduced muscle protein synthesis in the GL261 hosts, as evidenced by decreased puromycin incorporation, which also aligns with lower muscle IGF-1 expression levels, typically found in adult cancer cachexia models (113). Together, these mechanisms could explain, at least in part, the stunted muscle development observed, rather than more severe muscle loss seen in other cancer models.

Another major contributing factor to skeletal muscle atrophy was found to relate to mitochondrial dysfunction (114). Specifically, we and others have previously reported that cancer cachexia drives the dysregulation of mitochondrial homeostasis in adult mouse models (24, 26, 27, 52, 115), frequently accompanied by reductions in PGC-1α (116–121) and Mitofusin 2 levels (64, 118) and in line with abnormal mitochondria in skeletal muscle. PGC-1α has been described as the master regulator of mitochondrial biogenesis (122), and deficiency of this protein has been correlated with mitochondrial structural derangements, decreased insulin sensitivity, glucose tolerance, and reduced transcriptional activity (123, 124). Data from our group demonstrated that PGC-1α overexpression was sufficient to attenuate the decrease in mitochondrial cell signaling induced by a chemotherapy regimen, thus also preserving muscle mass and function in 2- and 18-month-old mice (125). Mitofusin 2, a downstream target of PGC-1α (126), is a marker of mitochondrial fusion (49) and modulates the tethering of the endoplasmic reticulum to neighboring mitochondria, ultimately governing calcium signaling. This tethering predisposes mitochondria to excessive calcium transfer, potentially leading to apoptosis and muscle loss (127, 128). Conversely, overexpression of Mitofusin 2 was shown to attenuate cancer-induced muscle wasting (129). Another abundant mitochondrial membrane protein, VDAC, regulates mitochondrial calcium flux and plays a role in apoptosis by transferring apoptotic calcium signals into the mitochondria (130, 131). Lower VDAC protein content normally suggests reduced mitochondrial content in cachectic tumor hosts (119, 132, 133), whereas increased VDAC permeability (134) can lead to increased calcium flux, calcium dysregulation, mitochondrial swelling (131) and the opening of permeability transition pores, altogether resulting in apoptosis (135, 136). Interestingly, in the present study we observed an upregulation of PGC-1α and Mitofusin 2 in the muscle of the GL261 hosts, which is in contrast with previous observations in the traditional cancer cachexia models. Similarly, the increase in VDAC expression observed in the pediatric tumor hosts is unique, as adult cancer cachexia models frequently show no changes (137) or decreases in VDAC protein expression in skeletal muscle (119, 133). Nonetheless, while one study described an increase in PGC-1α and Mitofusion 2 gene expression in mice bearing hepatomas (135), it is worth noting that in the context of cachexia, markers associated with positive changes in mitochondrial dynamics are typically downregulated; thus, we can speculate that our findings could be indicative of calcium dysregulation or a compensatory change in response to the tumor.

Overall, our observations are unique, especially since there is minimal research examining the muscle histological and functional consequences and the molecular mechanisms governing these effects in mouse models for the study of pediatric tumors. As such, our study may serve as foundational work to address these issues and contribute to the understanding of the mechanisms responsible for the onset of musculoskeletal detriments consequential to pediatric cancers. It is also worth noting that this research focused narrowly on the effects of glioblastoma tumors and only examined select mediators of stunted development, thus representing a limitation of our study. Another confounding variable was also the use of young mice that were in the exponential growth phase of their lifespan. Indeed, here we observed an increase in body weight of 39% in the control group, accompanied by a 65% gain (including the tumor weight) in the GL261 hosts. Despite this, we noted a reduction in puromycin incorporation in skeletal muscle, indicative of reduced protein anabolism, which is also typically observed in adult cancer models. However, in contrast to ‘conventional’ cachexia signaling, we also described a decrease in the gene expression of E3 muscle-specific ubiquitin ligases, thus representing a unicum. Moreover, given that mice between 4-7 weeks of age can be considered as ‘adolescent’ or ‘young adult’ (138), the age of the mice employed in the present study (i.e., 4 weeks) may be beyond the ideal ‘pediatric’ age window (<3 weeks). Due to logistical constraints, such as animal delivery timing and mandatory acclimation period at our animal facility, 4 weeks of age is the earliest possible start for this kind of study, although the use of animals born in house may allow by-passing such restrictions. Another limitation of the current research is the use of GL261 subcutaneous inoculations instead of orthotopic surgical implantations. Indeed, the use of orthotopic CNS cancers may also contribute to investigate the brain microenvironment, especially considering that the hypothalamus, a critical brain region for regulating metabolism and appetite, was found to be affected by tumor-derived factors, thus also disrupting normal neuroendocrine functions and promoting systemic consequences, as reviewed in (139). Lastly, in the present study we did not investigate how chemotherapeutics influence the musculoskeletal phenotype in the GL261 hosts. Indeed, our previous work shed light on the detrimental effects of anti-cancer agents on musculoskeletal health in adult animals (25, 52, 64, 140, 141). Recently we also demonstrated persistent musculoskeletal consequences in pediatric mice administered chemotherapeutic regimen (36). Interestingly, work by De Witt *et al*. reported greater reductions in body mass in mice bearing GL261 tumors in conjunction with chemotherapy, although the studies used older mice (3 months) and did not carefully characterize the effects on muscle mass (28). Future studies will investigate the mechanisms responsible for the systemic long-term effects of cancer and its therapies in pediatric patients and investigate potential therapeutic targets to attenuate the blunted muscle development.

In conclusion, our study suggests that glioblastoma tumors can induce detrimental effects, as demonstrated by the decrease in carcass weight, skeletal muscle mass, muscle cross sectional area, organ weight, and an unconventional cell signaling cascade. Overall, the present study demonstrates that growth of GL261 tumors contributes to the stunted development of young mice, as evidence by reduced muscle mass and muscle fiber size. These findings are supported by evidence that pro-cachexiogenic signatures normally observed in adult mouse models (e.g., heightened STAT3 phosphorylation, ubiquitination, autophagy, and reduced muscle protein synthesis) are similarly activated in the muscle of young GL261 hosts, whereas our unique finding of decreased E3 muscle-specific ubiquitin ligase expression may also contribute to explain the milder effects compared to other cancer models.

## Funding

This study was supported by the Department of Surgery and the Department of Otolaryngology – Head & Neck Surgery at Indiana University School of Medicine, the Department of Pathology and the Comprehensive Cancer Center at University of Colorado Anschutz Medical Campus, and by grants from NIH/NIAMS (R01AR079379, R01AR080051), American Cancer Society (132013-RSG-18-010-01-CCG), and Cancer League of Colorado, Inc. and MDC/Richmond American Homes Foundation (AWD-242527-AB) to AB. JRH, NAJ and CSC were supported by T32 Institutional Training Grants from NIH (AR065971, CA190216, AR080630).

## Acknowledgments

The #12G10 anti-Tubulin, MF-20, and #MANDRA11(8B11) monoclonal antibodies were obtained from the Developmental Studies Hybridoma Bank, created by the NICHD of the NIH and maintained at The University of Iowa, Department of Biology, Iowa City, IA. The murine GL261 cells were kindly provided by Dr. Reza Saadzadeh (Indiana University School of Medicine, Indianapolis, IN, USA)

---

**Figure S1. GL261 culture medium promote myotube atrophy.** Myotube diameter (expressed as µm) of C2C12 myotubes exposed to GL261 conditioned medium **(A)**. Representative images of control (Con) and myotubes exposed to GL261 CM **(B)**. Scale bar: 100 µm. Data expressed as mean ± SD. Significant differences: ***p<0.001.

**Figure S2.** Schematic of experimental study.

**Figure S3. GL261 tumors reduce gonadal fat and heart weight, and increase spleen weight with no effect on kidney and liver weights.** Gonadal fat (A), heart (B), spleen (C), liver (D), and kidney (E) weights were normalized to initial body weight (IBW) and expressed as weight/100 mg IBW. Data reported as mean ± SD. Significant differences: *p<0.05, **p<0.001.

**Figure S4. GL261 hosts exhibit changes in muscular force and alterations in neuromuscular function.** EDL *ex-vivo* force measured between 10-150Hz (A), EDL peak *ex vivo* force measured at 150Hz (B), and peak *in vivo* plantar flexion torque in control mice and GL261 hosts. *Ex vivo* force was expressed as kN/m^2^. *In vivo* torque was expressed as mN•m. Base-to-peak compound muscle action potential [CMAP (b-p); expressed in millivolts (mV)] (D), single motor unit potential [SMUP; expressed in microvolts (μV)] (E), and motor unit number estimation [MUNE; expressed as number of motor units] of the triceps surae muscle in control and GL261-bearig mice. Data expressed as mean ± SD. Significant differences: *p<0.05, **p<0.01.

**Figure S5. GL261 yielded higher interleukin-6 (IL-6) and no differences in IGF-1 plasma concentrations.** Values expressed as pg/mL. Data expressed as mean ± SD. Significant differences: **p<0.01.

## Notes

### Competing Interest Statement

The authors have declared no competing interest.

