## Supplementary figures and images for "GL261 glioblastoma induces delayed body weight gain and stunted skeletal muscle growth in young mice"

### Figure S1

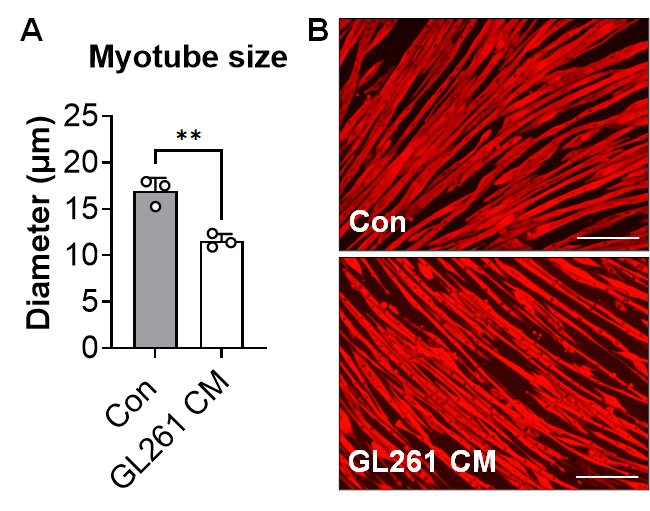

### Figure S2

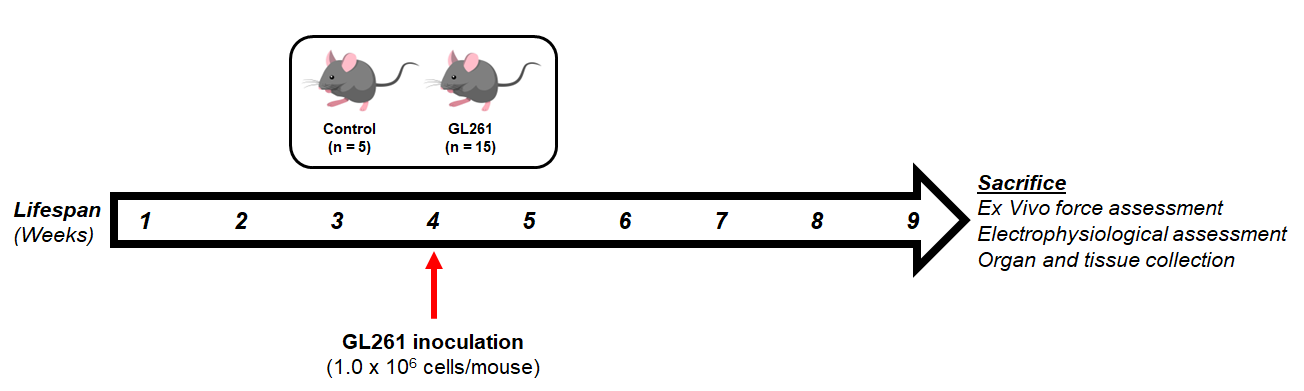

### Figure S3

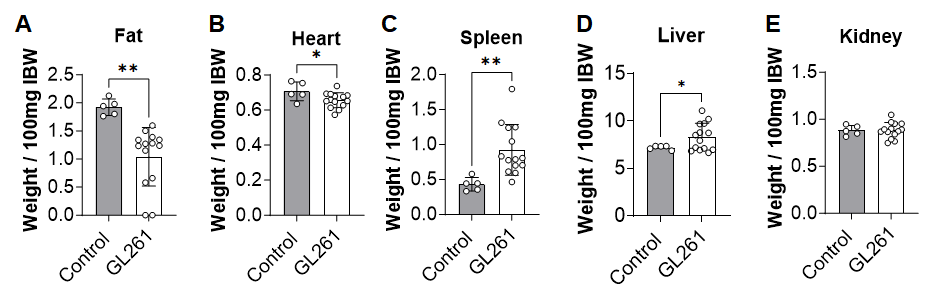

### Figure S4

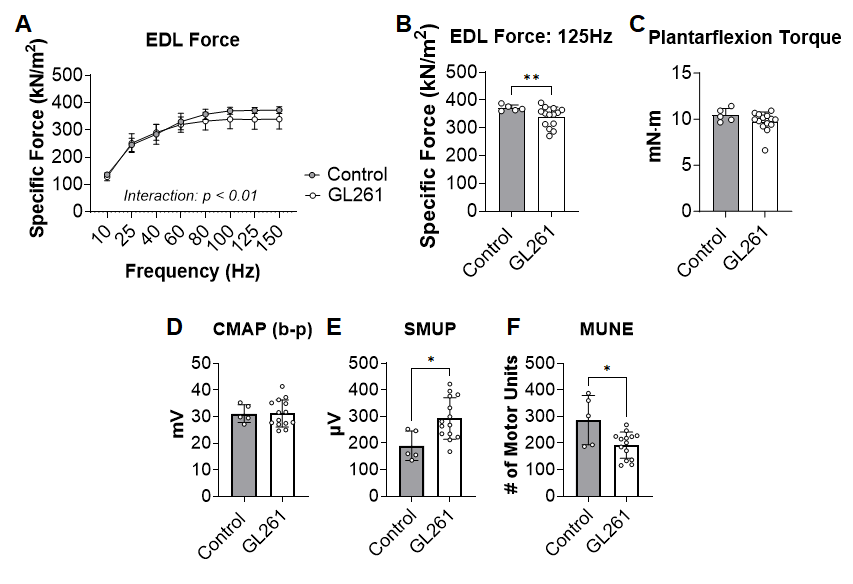

### Figure S5

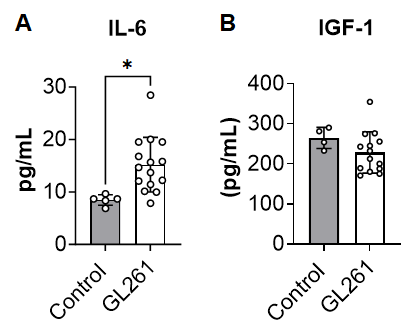
